# ESR2 regulates indian hedgehog signaling in neonatal rat ovary

**DOI:** 10.1101/2021.12.01.470808

**Authors:** Iman Dilower, V. Praveen Chakravarthi, Eun B. Lee, Subhra Ghosh, Shaon Borosha, Richita Roy, Saeed Masumi, Anohita Paul, Hindole Ghosh, Michael W. Wolfe, M. A. Karim Rumi

**Affiliations:** Pathology and Laboratory Medicine, University of Kansas Medical Center, Kansas City, KS 66160; Molecular and Integrative Physiology, University of Kansas Medical Center, Kansas City, KS 66160; Reproduction and Perinatal Research, University of Kansas Medical Center, Kansas City, KS 66160

**Keywords:** Ovary, *Esr2*, RNA-seq, *Ihh* and *Hhip*

## Abstract

The transcriptional regulatory function of estrogen receptor β (ESR2) is essential for the regulation of primordial follicle activation (PFA). Increased PFA due to the loss of ESR2 becomes evident as early as postnatal day 8 (PND8). To identify the ESR2-regulated genes that control PFA, we performed RNA-seq analyses of wildtype, and *Esr2* knockout (*Esr2*^*KO*^) neonatal rat ovaries collected on PND4, PND6, and PND8. Among the differentially expressed genes in *Esr2*^*KO*^ ovaries, indian hedgehog (*Ihh*) displayed the highest downregulation among the ovary enriched genes. IHH regulated genes including *Hhip* as well as the steroidogenic enzymes were also downregulated in *Esr2*^*KO*^ rat ovaries. Remarkably, the expression of *Ihh* in *Esr2*^*KO*^ ovaries was not upregulated despite the high levels of *Gdf9* and *Bmp15*, which are known regulators of *Ihh* expression in granulosa cells. Our findings suggest that indian hedgehog signaling in the neonatal rat ovary is dependent on ESR2.

## 1. Introduction

Estrogen receptor β (ESR2) is the predominant estrogen receptor in mammalian ovary and is essential for follicle development and ovulation (1-3). ESR2 also plays a gatekeeping role to control the rate of primordial follicle activation (PFA) (4). Loss of ESR2 increases PFA leading to premature ovarian insufficiency in *Esr2* knockout (*Esr2*^*KO*^) rats (4, 5). Regulation of PFA is a complex mechanism that involves a number of negative regulators within the PI3-kinase and mTOR pathways (6-8). Several transcriptional regulators like FOXO3A also act like ESR2 to inhibit induction of PFA (9). Although the loss of ESR2 augments the activation of AKT, ERK, and mTOR pathways in *Esr2*^*KO*^ ovaries, the exact molecular mechanism of ESR2-mediated regulation of PFA remains unclear (4).

During ovarian follicle development, activated granulosa cells (GCs) express hedgehog proteins, which are essential for the development, growth, and differentiation of theca cells (TCs) (10-16). The loss of either hedgehog signaling, or its aberrant activation leads to defective differentiation of TCs and failure of ovulation (12, 15). The indian hedgehog (*Ihh*), and the desert hedgehog (*Dhh*) genes are expressed in GCs, whereas the hedgehog receptor *Ptch1/Ptch2*, the signal transducer *Smo*, as well as the hedgehog target genes like *Gli1/Gli2/Gli3* transcription factors are expressed in TCs (17, 18). It is also important to note that ESR2 is predominantly expressed in the GCs, while ESR1 is expressed in TCs (19, 20). However, it remains unknown if ESR2 plays any regulatory role of hedgehog gene expression in GCs or ESR1 on the expression of hedgehog receptors.

We previously identified that the canonical transcription function of ESR2 is responsible for the regulation of PFA (4). In this study, we have explored the ESR2-regulated genes in developing rat ovaries to understand the mechanisms underlying ESR2 regulation of PFA. We identified that the expression *Ihh* and its downstream target *Hhip* is completely dependent on the presence of ESR2 in neonatal rat ovaries.

## 2. Results

### 2.1. Alerted follicle activation in Esr2^KO^ rats

We detected an increased activation of PdFs in *Esr2*^*KO*^ rat ovaries compared to age-matched wildtype. Increased number of PrFs were clearly evident on PND8 *Esr2*^*KO*^ rat ovaries (**Fig.1 E-I**). While the cortical region in wildtype rat ovaries predominantly contained PdFs, it was replaced by PrFs in *Esr2*^*KO*^ rat ovaries. The total follicular counts of ovarian follicles in various stages of development remained similar between wildtype and *Esr2*^*KO*^ rat ovaries (**Fig.1I**).

**Figure 1.**
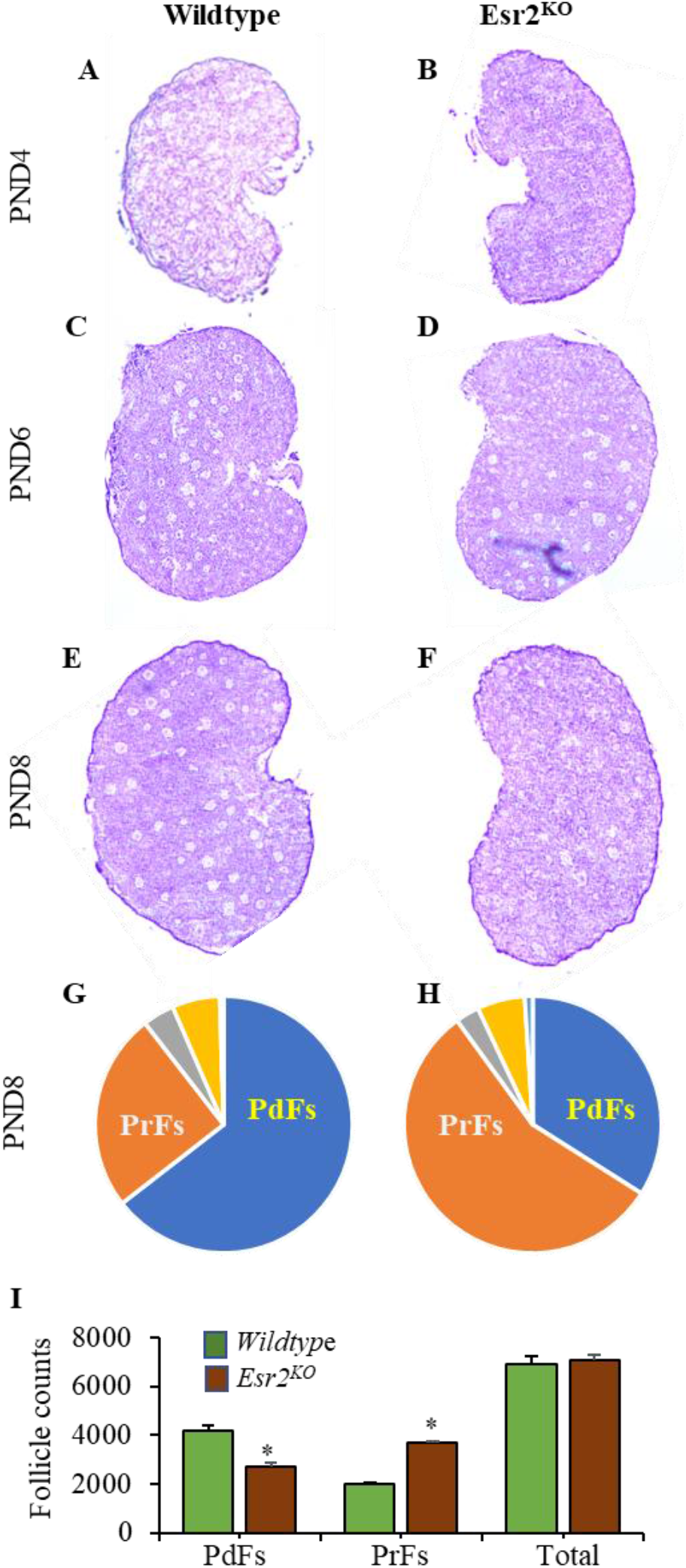
Increased PFA in *Esr2*^*KO*^ ovaries. Hematoxylin and eosin-stained sections of wildtype and *Esr2*^*KO*^ PND4 (**A, B**), PND6 (**C, D**), and PND8 (**E, F**) rat ovaries showed increased activation of PdFs in *Esr2*^*KO*^ rats. Follicle counting in PND8 wildtype and *Esr2*^*KO*^ rat ovaries showed significantly decreased numbers of PdFs and increased numbers of PrFs (**G-I**). However, the total count of ovarian follicles in wildtype and *Esr2*^*KO*^ ovaries remained similar (I). Data shown as mean ± SE, ^*^ *p*<0.05, n ≥ 3.

### 2.2. ESR2-regulated genes in neonatal rat ovaries

To identify the transcriptional targets of ESR2, we performed RNA-sequencing of PND4, PND6 and PND8 *Esr2*^*KO*^ and age matched wildtype rat ovaries. Analyses of the RNA-seq data identified a large number of differentially expressed genes (DEGs) in *Esr2*^*KO*^. For further analyses we focused on the DEGs related to hedgehog, steroidogenesis, and folliculogenesis pathways (**Fig. 2 A-C**).

**Figure 2.**
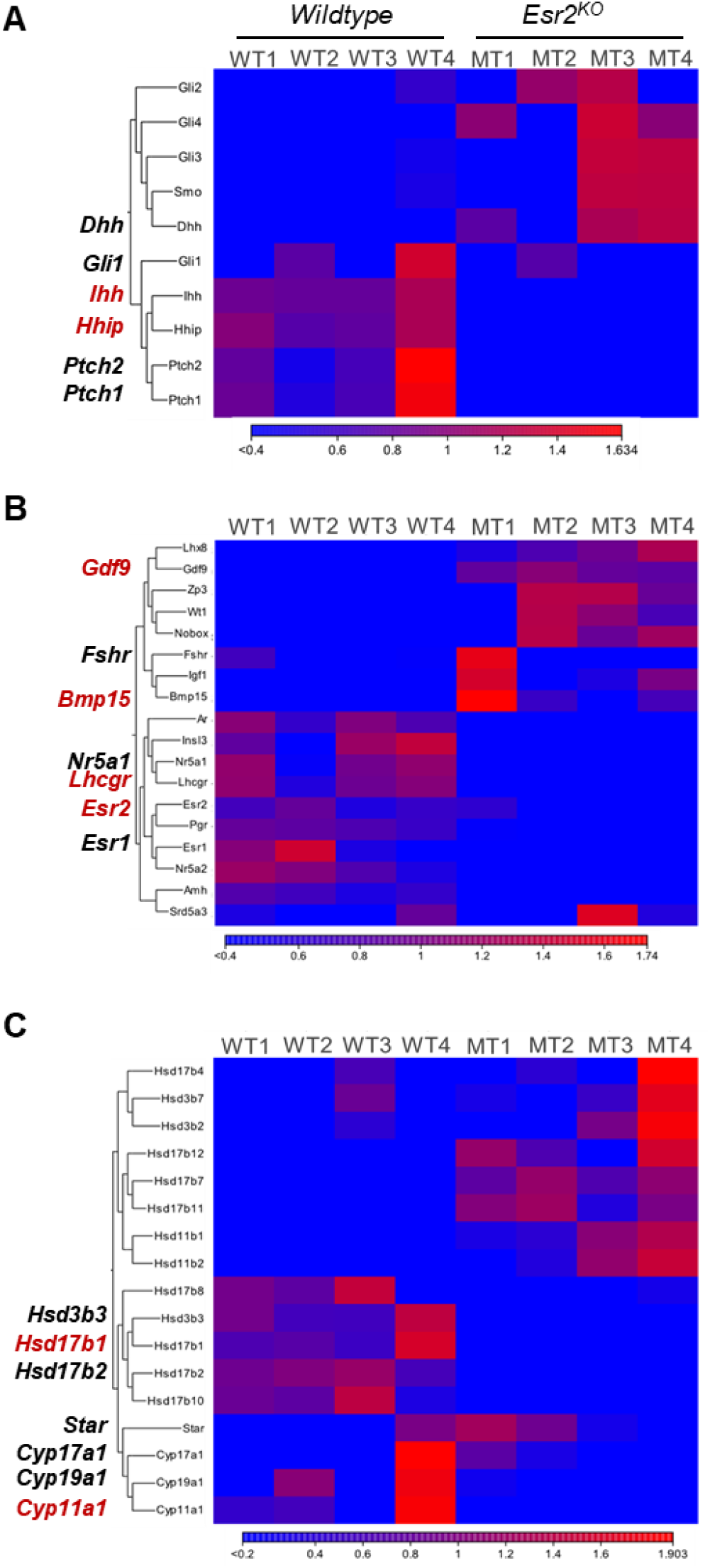
RNA-seq analysis of PND8 rat ovaries. RNA-seq analysis of PND8 rat ovaries showing differential expression of genes related to hedgehog pathway (A), steroidogenesis (B) and the key genes involved in folliculogenesis (C).

### 2.3. Disruption of indian hedgehog signaling

Among the transcripts with TPM values ≥1, *Ihh* (**Fig. 3A**) showed the highest downregulation. Next to *Ihh* was the downregulation of IHH target gene *Hhip* (**Fig. 3B**).

**Figure 3.**
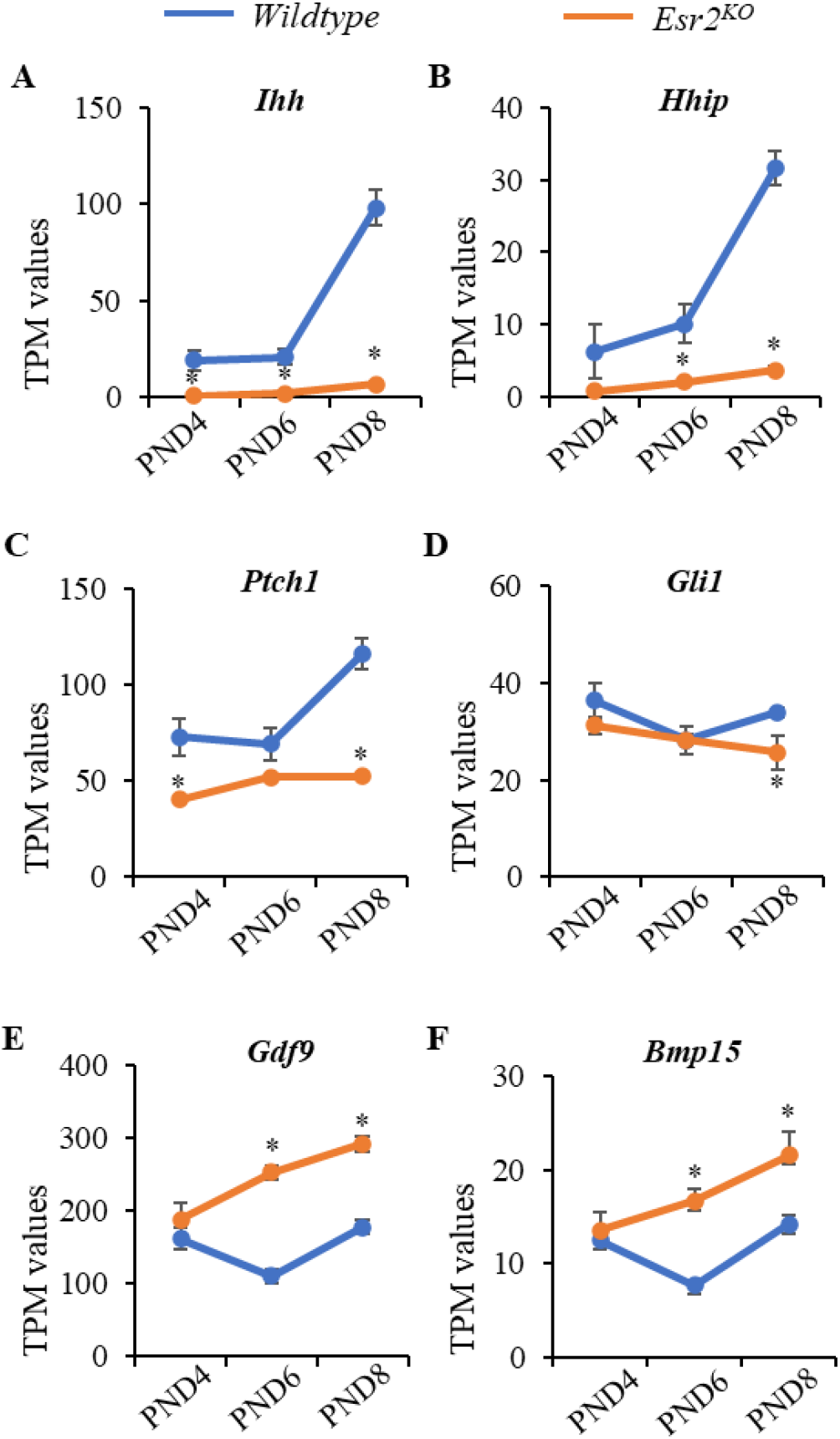
Loss of ESR2 disrupts indian hedgehog signaling. RNA sequencing analysis of PND4, PND6, and PND8 showed marked downregulation of *Ihh*, its downstream targets *Hhip* and *Gli1*, as well as its receptor *Ptch1* (**A-D**) in *Esr2*^*KO*^ ovaries. However, its known regulators *Gdf9* (**E**) and *Bmp15* (**F**) were significantly upregulated in *Esr2*^*KO*^ ovaries. Data shown as mean±SE TPM values, ^*^ *p*≤ 0.05, n=4.

Downregulation of both *Ihh* and *Hhip* was consistent and increased from PND4 to PND8 (p=0.00) (**Fig. 3A, B**).

Expression of hedgehog receptor *Ptch1* and hedgehog target *Gli1* was also downregulated in *Esr2*^*KO*^ rat ovaries (**Fig. 3C, D**). However, the expression of *Gdf9* and *Bmp15* was significantly upregulated in *Esr2*^*KO*^ rat ovaries (**Fig. 3E, F**). The RNA-seq data were verified by RT-qPCR, which showed similar extent of *Ihh, Hhip, Ptch1, Gli1* downregulation in PND8 *Esr2*^*KO*^ rat ovaries (**Fig. 4A-D**). Upregulation of *Gdf9* and *Bmp15* expression was also evident in PND8 *Esr2*^*KO*^ rat ovaries (**Fig. 4E, F**).

**Figure 4.**
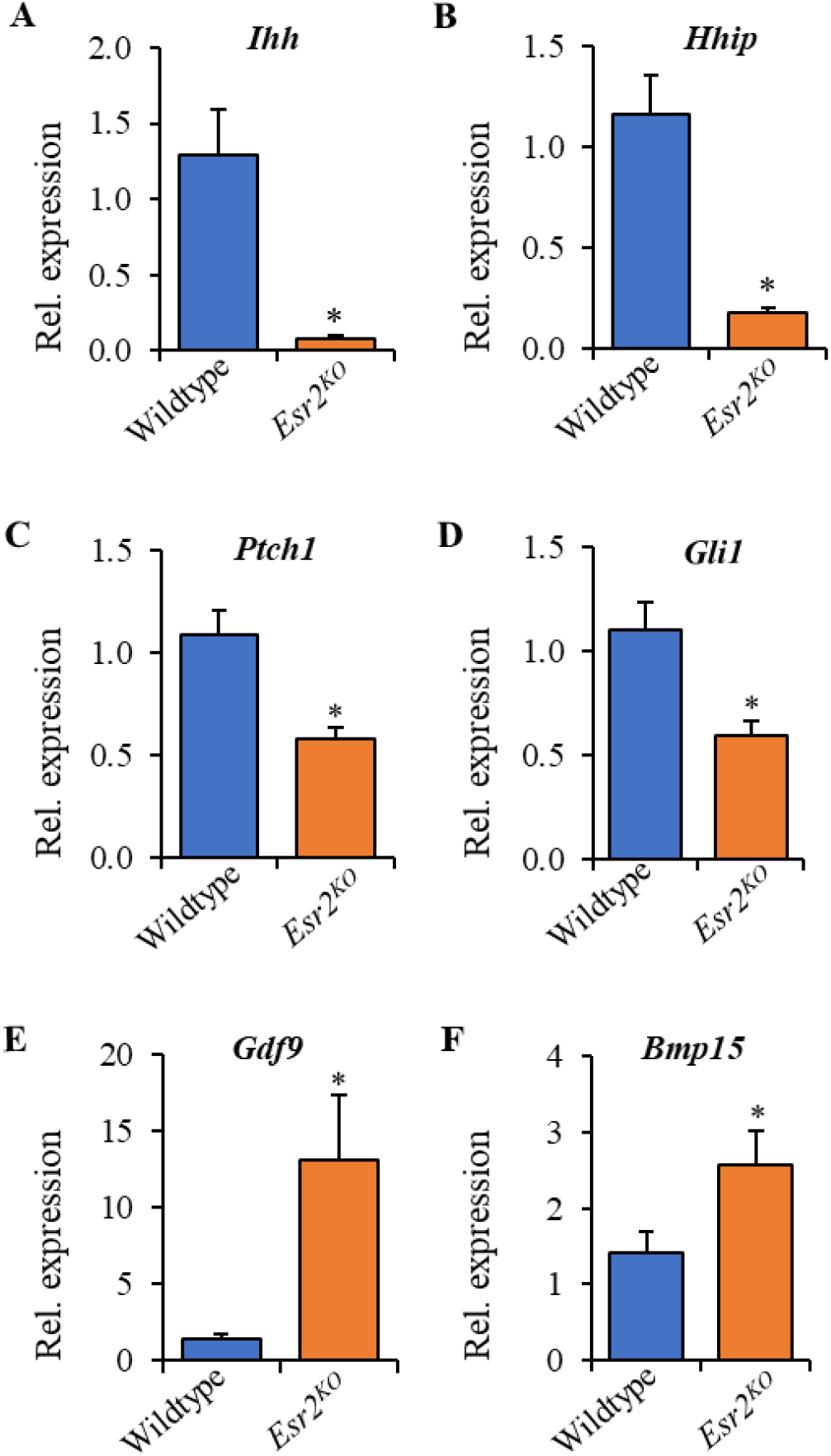
Verification of hedgehog signaling genes. RNA-seq data were verified by RT-qPCR on PND8 *Esr2*^*KO*^ rat ovaries. RT-qPCR confirmed downregulation of *Ihh, Hhip, Ptch1*, and *Gli1* (**A-D**), despite the marked upregulation of *Gdf9* and *Bmp15* (**E, F**). RT-qPCR data are shown as mean ± SE. ^*^*p* < 0.05, n = 6.

### 2.4. Altered expression of steroidogenic enzymes

IHH signaling plays an important role in regulating the expression of steroidogenic enzymes. While analyzing the RNA-seq data, we identified that loss of ESR2 in *Esr2*^*KO*^ rat ovaries disrupted the expression of *Hsd17b1* and *Cyp11a1* (**Fig. 5A, C**). Expression of gonadotropin receptor *Lhcgr* was significantly downregulated in *Esr2*^*KO*^ rat ovaries (**Fig. 5F**). However, the expression of *Fshr* and FSH responsive gene *Star* remained unchanged in *Esr2*^*KO*^ rat ovaries (**Fig. 5B, E**).

**Figure 5.**
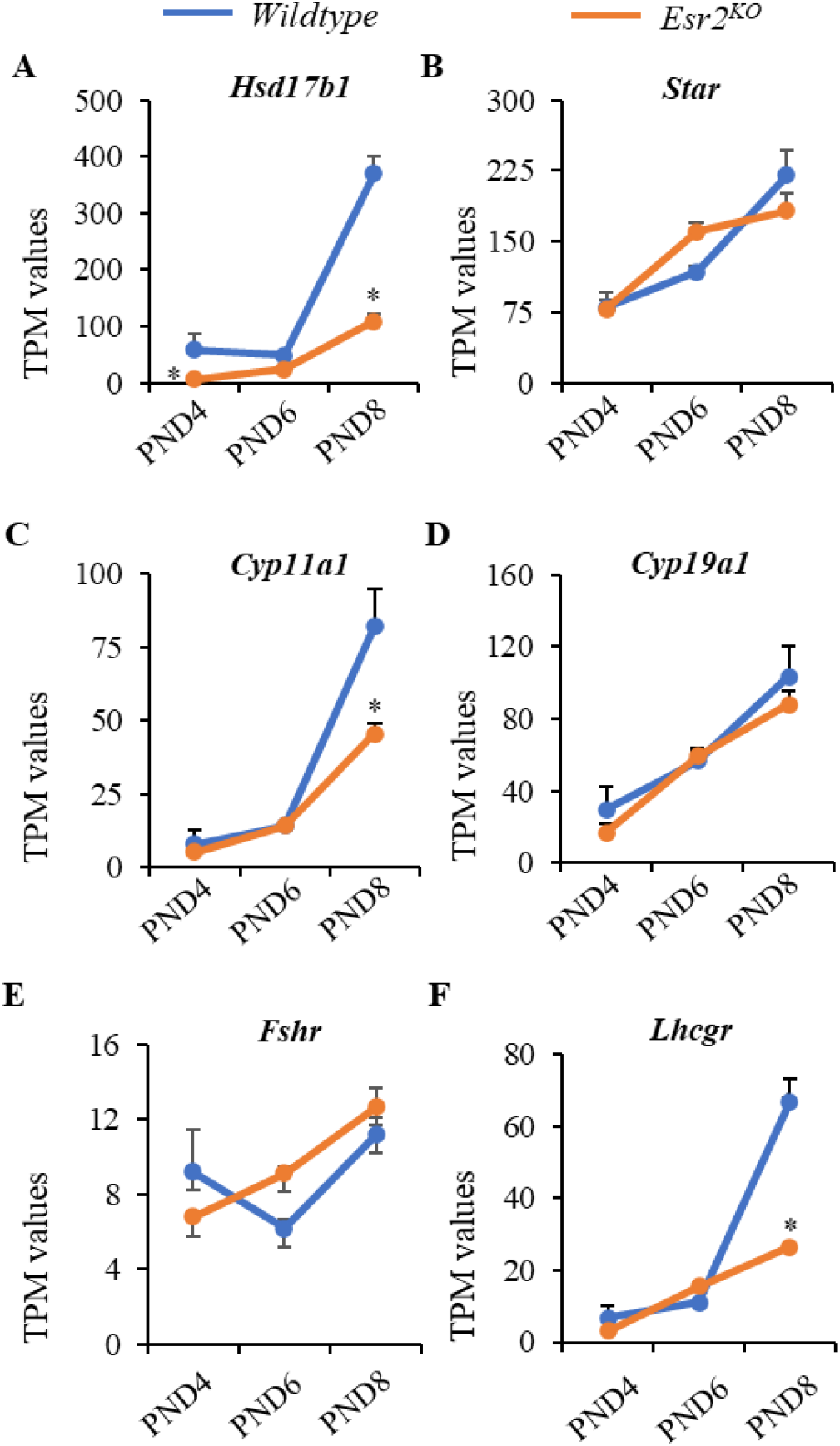
Loss of ESR2 disrupts expression of steroidogenic enzymes. RNA-seq analysis of PND4, PND6, and PND8 showed marked downregulation of *Hsd17b1*, and *Cyp11a1* (**A, C**) in *Esr2*^*KO*^ ovaries. However, the expression of *Star*, and *Cyp19a1* did not show any significant changes in *Esr2*^*KO*^ ovaries (**B, D**). While *Fshr* remained unchanged, expression of *Lhcgr* was downregulated in PND8 (E, F). Data shown as mean±SE TPM values, ^*^*p*≤ 0.05, n=4.

The RNA-seq data were verified by RT-qPCR, which showed similar extent of *Hsd17b1* and *Cyp11a1* downregulation in PND8 *Esr2*^*KO*^ rat ovaries (**Fig. 6A,C**). Downregulation of *Lhcgr* expression was also evident in PND8 *Esr2*^*KO*^ rat ovaries (**Fig. 6F**).

**Figure 6.**
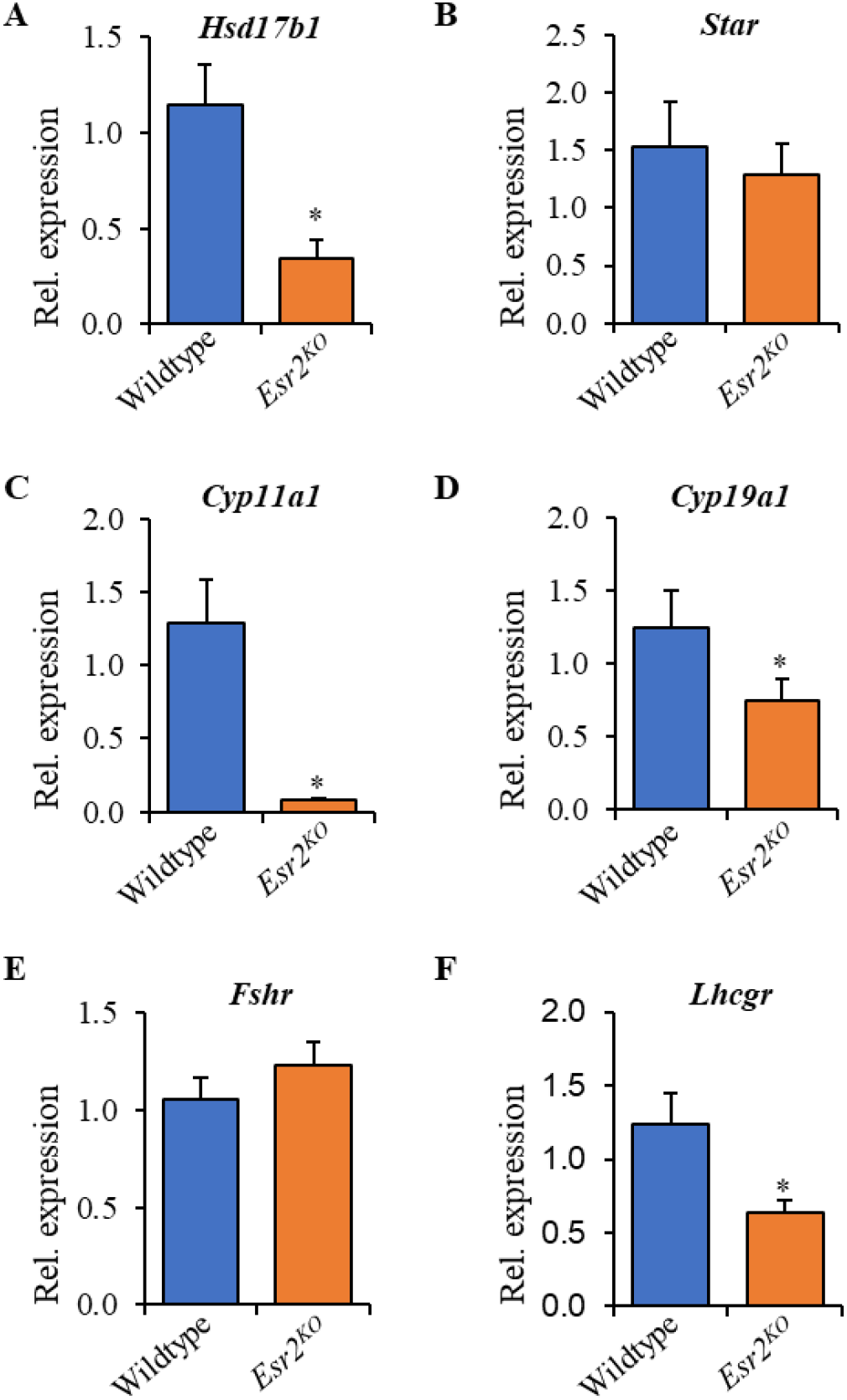
Verification of the expression of steroidogenic genes. RNA-seq data were verified by RT-qPCR on PND8 *Esr2*^*KO*^ rat ovaries. RT-qPCR analyses confirmed the downregulation of steroidogenic enzymes *Hsd17b1, Cyp11a1* and *Cyp19a1*. Downregulation of gonadotropin receptor *Lhcgr* was also similar to RNA-seq data. RT-qPCR data are shown as mean ± SE. ^*^*p* < 0.05, n = 6.

## 3. Discussion

Findings of this study suggest an essential role of ESR2 signaling in the regulation of *Ihh* expression in neonatal rat ovary. GCs possess a high level of ESR2 expression (19, 20); thus, it is likely that *Ihh* expression in GCs is regulated by this ligand activated transcription factor. GCs in the activated ovarian follicles express the *Ihh* gene, which in turn act on premature TCs for induction of their development and differentiation (12, 21). Loss of *Ihh* expression in neonatal rat ovary was associated with a loss of *Hhip* expression in a similar fashion that was reported in *Ihh*^*KO*^ mouse ovary (13).

In addition to the development and differentiation of TCs, hedgehog signaling, particularly IHH, plays a pivotal role on regulation of ovarian steroidogenesis (14). In this study, we observed that lack of ESR2 dysregulated the expression of steroidogenic enzymes in *Esr2*^*KO*^ rat ovaries. Previous studies in *Ihh*^*KO*^ mice also reported a similar impact of lack of *Ihh* expression on ovarian steroidogenesis (13).

RNA-seq analyses identified altered expression of multiple steroidogenic enzymes, which can be either of GC or TC origin. It is well-known that loss of ESR2 affects the expression of GC-genes involved in steroidogeneses including *Cyp11a1* and *Cyp19a1* (1, 2). We have observed that *Ihh* target gene *Hhip*, which is of TC origin, is dramatically downregulated in *Esr2*^*KO*^ rat ovaries. Thus, it is likely to observe downregulation of the steroidogenic enzyme genes in TCs, particularly those that are regulated by IHH signaling.

Previous studies have suggested that GDF9 and BMP15 expressed by the oocytes in activated follicles induce the expression of *Ihh* in mouse GCs (12, 22). In this study, we observed a very low level of *Ihh* and *Hhip* gene expression in neonatal *Esr2*^*KO*^ rat ovaries despite high levels of *Gdf9* or *Bmp15* expression. These observations indicate that GDF9 or BMP15 requires the presence of transcriptional regulator ESR2 to induce *Ihh* expression in GCs.

Although mice harboring ovary-specific *Ihh*^*KO*^ did not exhibit any ovulatory dysfunction (13), it does not exclude a possible role of IHH signaling in the regulation of PFA. Moreover, the representative images of *Ihh*^*KO*^ mouse ovaries appeared smaller than that of wildtype mouse ovaries (13), which is comparable to the relatively smaller sizes of *Esr2*^*KO*^ mouse or *Esr2*^*KO*^ rat ovaries (1, 23). We hypothesize that ESR2-regulated expression of *Ihh* from the activated follicles may act on dormant PdFs to control excessive PFA, which is observed in *Esr2*^*KO*^ rat ovaries (4). However, further studies are required to prove a mechanistic role of IHH in the regulation of PFA.

## 4. Materials and methods

### 4.1. Animal models

*Esr2*^*KO*^ mutant rats were included in this study. *Esr2*^*KO*^ rats were generated by targeted deletion of exon 3 in *Esr2* gene (1). Experimental animals were generated by breeding heterozygous mutant male and female rats carrying the *Esr2*^*KO*^ alleles. Rats were screened for the presence of mutations by PCR using tail-tip DNA samples (RED extract-N-Amp Tissue PCR Kit, Millipore-Sigma) as described previously (1, 24). Ovaries were collected from 4-to-8-day old *Esr2*^*KO*^ and age-matched wildtype rats for histological, and molecular studies. All animal experiments were performed in accordance with the protocols approved by the University of Kansas Medical Center Animal Care and Use Committee.

### 4.2. Histological evaluation of ovarian phenotypes

Ovaries were collected from 4-to-8-day old *Esr2*^*KO*^ and age-matched wildtype rats. One ovary from each rat was embedded in OCT, frozen immediately and preserved at - 80°C freezer till sectioning. The other ovary was snap-frozen in liquid nitrogen and preserved at -80°C freezer till processed for RNA extraction. Histological sections were prepared using a cryotome. The frozen sections were prepared from whole ovaries at 6 µm thickness and placed on charged glass slides (Fisher Scientific). Ovary sections were stained with hematoxylin and eosin (H&E) following standard procedure (25). H&E-stained sections were thoroughly examined for follicle morphology and counted for follicles in each stage of development as we have described previously (4).

### 4.3. Detection of differentially expressed genes

Gene expression at the mRNA level was evaluated by RNA sequencing (RNA-seq), conventional RT-PCR, and RT-qPCR analyses. Total RNA was extracted from the whole ovary with TRI Reagent (Millipore-Sigma) and those with RIN values ≥ 9 were considered for library preparation. 500 ng of total RNA from each sample was used for the RNA-seq library preparation using the TruSeq stranded mRNA kit (Illumina, San Diego, CA) following the manufacturer’s instructions. The cDNA libraries were evaluated for quality at the KUMC Genomics Core and then sequenced on an Illumina HiSeq X sequencer at Novogene Corporation, Sacramento, CA. All RNA-seq data have been submitted to the Sequencing Read Archive (SRX6955095-6955104). RNA-seq data were analyzed using CLC Genomics Workbench

### 4.4. Validation of the differentially expressed genes

Total RNA was extracted from the PND 8 ovaries using TRI Reagents (Sigma-Aldrich). 1000 ng of total RNA from each sample was used for the preparation of cDNAs in 20µl volume using High-Capacity cDNA Reverse Transcription Kits (Applied Biosystems, Foster City, CA). RT-qPCR amplification of cDNAs was carried out in a 10µl reaction mixture containing Applied Biosystems Power SYBR Green PCR Master Mix (ThermoFisher Scientific). Amplification and fluorescence detection of RT-qPCR were carried out on Applied Biosystems QuantStudio Flex 7 Real Time PCR System (ThermoFisher Scientific). The ΔΔCT method was used for relative quantification of target cDNA abundance representing mRNA expression level normalized to Rn18s (18S rRNA).

### 4.5. Statistical analysis

Each RNA-seq library was prepared using pooled RNA samples from 3 individual wildtype or *Esr2*^*KO*^ rats. Each group of RNA-sequencing data consisted of four different libraries. For other experiments, each group consisted ovaries from a minimum of 6 rats, and all procedures were repeated for reproducibility. The data are presented as the mean ± standard error (SE). The results were analyzed using one-way ANOVA, and the significance of mean differences was determined by Duncan’s post hoc test, with *p*≤ 0.05. All the statistical calculations were done using SPSS 22 (IBM, Armonk, NY).

## Acknowledgments

This study was supported by funding from the KUMC SOM, COBRE, and K-INBRE.

